# SaProt: Protein Language Modeling with Structure-aware Vocabulary

**DOI:** 10.1101/2023.10.01.560349

**Authors:** Jin Su, Chenchen Han, Yuyang Zhou, Junjie Shan, Xibin Zhou, Fajie Yuan

## Abstract

Large-scale protein language models (PLMs), such as the ESM family, have achieved remarkable performance in various downstream tasks related to protein structure and function by undergoing unsupervised training on residue sequences. They have become essential tools for researchers and practitioners in biology. However, a limitation of vanilla PLMs is their lack of *explicit* consideration for protein structure information, which suggests the potential for further improvement. Motivated by this, we introduce the concept of a “<u>s</u>tructure-<u>a</u>ware vocabulary” that integrates residue tokens with structure tokens. The structure tokens are derived by encoding the 3D structure of proteins using Foldseek. We then propose SaProt, a large-scale general-purpose PLM trained on an extensive dataset comprising approximately 40 million protein sequences and structures. Through extensive evaluation, our SaProt model surpasses well-established and renowned baselines across 10 significant downstream tasks, demonstrating its exceptional capacity and broad applicability. We have made the code^1^, pre-trained model, and all relevant materials available at https://github.com/westlake-repl/SaProt.

## 1 Introduction

Proteins are fundamental to biological functions, and understanding them opens promising avenues in medical, pharmaceutical, and genetic research. Protein Language Models (PLMs), drawing inspiration from NLP methodologies, have emerged as the pivotal technology for representing proteins (Rao et al., 2019). Through self-supervised training on vast amounts of protein 1D residue sequences, PLMs have proven highly proficient in capturing long-range residue correlations, i.e., co-evolution (Anishchenko et al., 2017; Rao et al., 2020). Moreover, prominent PLMs like UniRep (Alley et al., 2019), ProtTrans (Elnaggar et al., 2021), ESM (Rives et al., 2019; Meier et al., 2021; Rao et al., 2021; Lin et al., 2022), and Evoformer (Hu et al., 2022; Jumper et al., 2021) have showcased outstanding performance across a diverse array of tasks pertaining to protein structure and function.

Despite the success of residue sequence-based pre-training, there’s a growing interest in utilizing protein 3D structures as training data, given their direct relevance to functions. Some work have demonstrated the potential of pre-training on experimentally determined protein structures (Yang et al., 2022; Hermosilla & Ropinski, 2022), but they are limited by the smaller number of highly accurate structures compared to residue sequences. Meanwhile, the breakthrough achieved by AlphaFold2 (AF2) (Jumper et al., 2021) in protein structure prediction has resulted in a substantial repository of structure data (Varadi et al., 2021), thus igniting interests in utilizing large-scale protein structures for training PLMs.

Currently, the development of structure-based PLMs based on large-scale predicted structures is still in an early stage, and existing research has certain limitations. For instance, well-known models like GearNet (Zhang et al., 2023b), still depend on a limited set of protein structures, utilizing around 800 thousand predicted structures from AF2. On the other hand, models like ESM-IF (Hsu et al., 2022) focus exclusively on specific protein tasks, such as protein inverse folding, rather than aiming for broader and more general-purpose representations.

In this paper, we aim to contribute to the biological community by introducing a large and more powerful PLM trained on extensive protein sequence and structure data. To achieve this, we introduce a “structure-aware (*SA*) vocabulary” that encompasses both residue and structure information of proteins. Specifically, we can employ vector quantization techniques (Van Den Oord et al., 2017) to discretize protein structures into 3D tokens. These tokens, similar in format to residue tokens, capture the geometric conformation information of each residue in relation to its spatial neighbors. Here, we simply utilize Foldseek (van Kempen et al., 2023), a purpose-built tool. Then, by combining the 3D tokens with residue tokens, we devise a *very intuitive yet innovative* vocabulary termed the *SA* alphabet. This enables the conversion of the original residue sequence into an *SA*-token sequence, serving as the input for existing residue-based PLMs. Through unsupervised training on massive protein *SA*-token sequences, we obtain a **S**tructure-**a**ware **Prot**ein language model named SaProt. To assess its performance, we comprehensively evaluate its capabilities across 10 widely recognized protein tasks. These tasks encompass a broad range of applications, including clinical disease variant prediction (Frazer et al., 2021), fitness landscape prediction (Dallago et al., 2021; He et al., 2024), protein-protein interaction (Nooren & Thornton, 2003), as well as diverse protein function predictions (Bileschi et al., 2022; Yu et al., 2023). To summarize, our main contributions are as follows:

- We introduce a structure-aware vocabulary that combines residue and 3D geometric feature for proteins. With the utilization of “*SA*” tokens, proteins, encompassing both primary and tertiary structures, can be effectively represented as a sequence of these novel tokens. The sequential nature, rather than the graph^2^ structure, of protein representation allows for seamless integration with advances in large-scale foundation AI models, such as BERT (Devlin et al., 2018), BART (Lewis et al., 2019), GPT (Brown et al., 2020), etc.
- By utilizing the *SA*-token protein sequence as input, we train a structure-enhanced PLM using the ESM (Lin et al., 2022) backbone as a case study, called SaProt. To our knowledge, SaProt stands out as the PLM currently trained with the largest number of protein structures, containing 650 million parameters. Its training lasted 3 months and utilized 64 NVIDIA 80G A100 GPUs, with a computational cost similar to ESM-1b (Rives et al., 2019).
- We evaluate SaProt across 10 renowned biological tasks. SaProt consistently exhibited improved performance compared to strong baselines, particularly models from the ESM family, including ESM-1b, ESM-1v (Meier et al., 2021), and ESM-2 (Lin et al., 2022), which are considered leading PLMs in the field.
- We conduct a series of enlightening ablation studies, unveiling previously unknown findings. One such finding is the potential overfitting issues that may arise when training PLMs by integrating predicted structures with BERT-style training. This discovery highlights a crucial consideration in the design of protein structure-based PLMs. Additionally, our experimental section sheds light on several intriguing observations through dissecting SaProt.

Additionally, we have made our code, model weight, and the associated datasets openly available. These materials are expected to be valuable for both the computational and biological communities.

## 2 Related Work

### 2.1 Residue Sequence-based Pre-training

Sequence-based pre-training methods treat protein residue sequences as natural language, enabling comprehensive representations via masked language modeling (MLM) (Devlin et al., 2018). Formally, a protein sequence is denoted as *P* = (*s*_1_, *s*_2_, …, *s*_*n*_), where *s*_*i*_ is a residue at the *i*_*th*_ position and *n* is the sequence length. During pre-training, a set of residues are randomly masked, resulting in the modified sequence *P*_*mask*_ = (*s*_1_, *< MASK >*, …, *s*_*n*_). The training objective is to predict masked residues by capturing dependencies between masked positions and surrounding context.

Residue-based PLMs have shown potential in generating universal protein representations. Rives et al. (2019), Heinzinger et al. (2019) and Vig et al. (2020) substantiate the ability of PLMs to predict protein structures and functions, while Rao et al. (2021) enhances capabilities via training on Multiple Sequence Alignment (MSA) data. For mutational effect prediction, Meier et al. (2021) and He et al. (2024) adopt ESM-1v for zero-shot prediction, and Notin et al. (2022) incorporate MSA as supplementary signals. Additionally, Lin et al. (2022), Chowdhury et al. (2022) and Wu et al. (2022b) predict protein structures from single sequences by applying large PLMs.

### 2.2 Structure-based Pre-training

Protein structure governs its function. The release of 200 million protein structures in AlphaFoldDB (Varadi et al., 2022) in July 2022 enables the construction of large-scale protein structure models. Protein structures are usually represented as graphs, denoted by 𝒢 = (𝒱, *ℰ*), with 𝒱 representing the set of *N* residues and ℰ representing the set of edges connecting the residues. These edges are typically based on the *Cα* distances between the residues. GNNs utilize 𝒢 for diverse pretraining strategies like contrastive learning (Hermosilla & Ropinski, 2022; Zhang et al., 2023b;a), self-prediction (Yang et al., 2022; Chen et al., 2023) and denoising score matching (Guo et al., 2022; Wu et al., 2022a). Another way inspired by AF2 involves incorporating structure features as contact biases into the attention maps within the self-attention module, e.g., Uni-Mol (Zhou et al., 2023).

However, the above structure-based models rely on either real structures from the Protein Data Bank (PDB) or a limited number of predicted AF2 structures. To the best of our knowledge, there are currently no “general-purpose” PLMs based on a large-scale set of predicted structures.

### 2.3 Foldseek

The initial goal of Foldseek (van Kempen et al., 2022) is to facilitate fast and accurate protein structure searches. To achieve this, Foldseek employs a VQ-VAE model (Van Den Oord et al., 2017) for encoding protein structures into informative tokens. These tokens, derived from 20 distinct 3Di states, are represented as *P* = (*f*_1_, *f*_2_, …, *f*_*n*_), where *f*_*i*_ represents the structure token at the *i*_*th*_ position and *n* is the sequence length. Foldseek achieves this encoding by identifying nearest neighbors and extracting features for individual residues.

A preprint by Heinzinger et al. (2023) introduces ProstT5, which enables bidirectional conversion between residue and Foldseek token sequences. ProstT5 excels at tasks like remote homology detection and protein design. However, it is not considered a general-purpose PLM (see Appendix A).

## 3 Idea of New Vocabulary

### 3.1 Preliminary Analysis

The goal of this paper is to develop a general-purpose PLM by leveraging predicted protein structures to serve multiple protein prediction tasks. Contrastive learning (CL) and BERT-style MLM training are currently two most prevalent pre-training approaches. However, CL primarily emphasizes on protein-level representation learning and performs poorly at the residue-level task. For instance, GearNet (Zhang et al., 2023b) and 3D-PLM (Hermosilla & Ropinski, 2022) trained by CL are not directly useful for predicting effects of amino acid mutations (Frazer et al., 2021).

We initially explored two intuitive approaches for protein structure modeling. The first approach involves treating the predicted structures from AF2 as a graph and employing GNNs for modeling, following (Yang et al., 2022)^3^. The second approach is to extract the distance and angle information between pairwise residues from structures, incorporating it as a structure bias in a Transformer (Vaswani et al., 2017) attention map. This approach was applied by Uni-Mol, ESMFold (Lin et al., 2022) and Evoformer. We evaluate the two models^4^ using the MLM objective as it can support both protein-level and residue-level tasks. It should be noted that the structure model in Uni-Mol, ESMFold, and Evoformer were initially designed for specific tasks with different loss functions, rather than being intended as general-purpose PLM. Therefore, it remains uncertain whether these neural networks would be effective when trained with predicted structures using the MLM objective.

Through two exploratory experiments, we noted that training directly using predicted structures yielded poor performance on the validation set containing real PDB structures (Figure 2). The decrease in loss on predicted structures did not correspond to a decrease in loss on real structures. This mismatch may be due to the fact that PLM has detected traces of AF2 predictions. Furthermore, inferior results were reported in downstream tasks (Table 8). Despite a substantial loss decrease on training data, these models failed to learn meaningful representations for downstream protein tasks.

**Figure 1:**
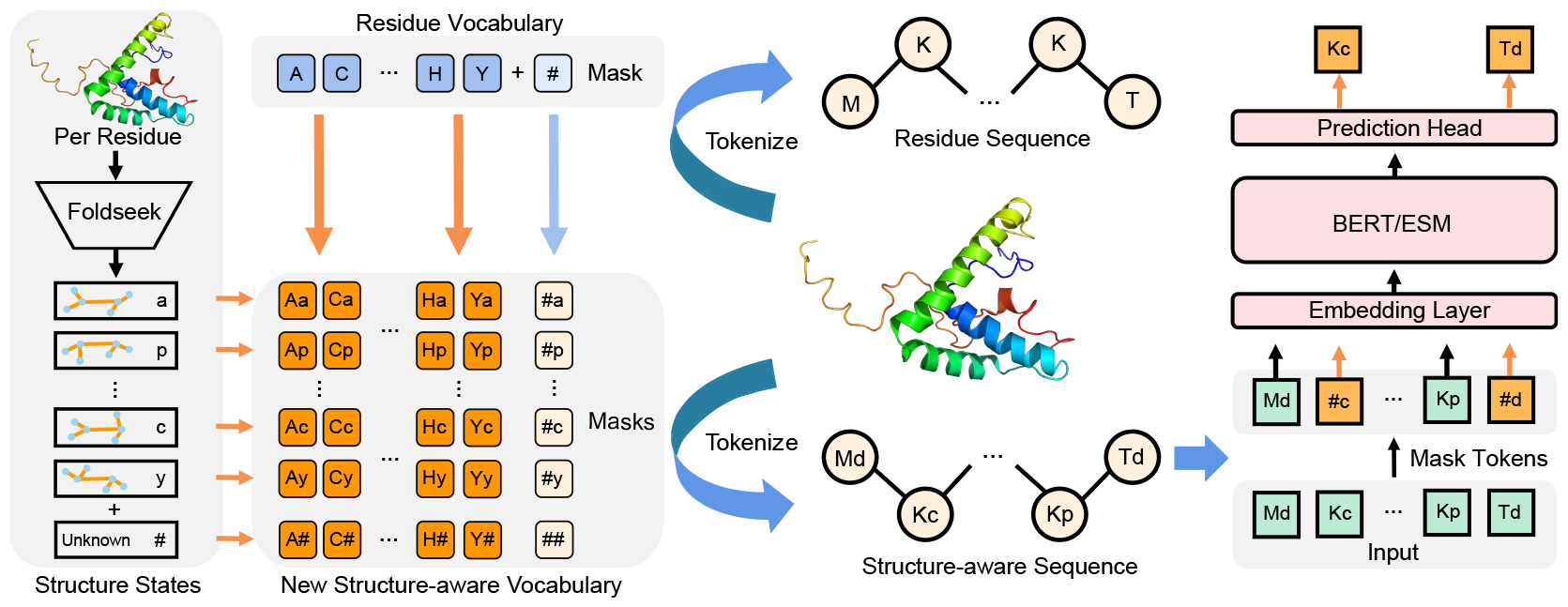
Framework of SaProt with the structure-aware vocabulary

**Figure 2:**
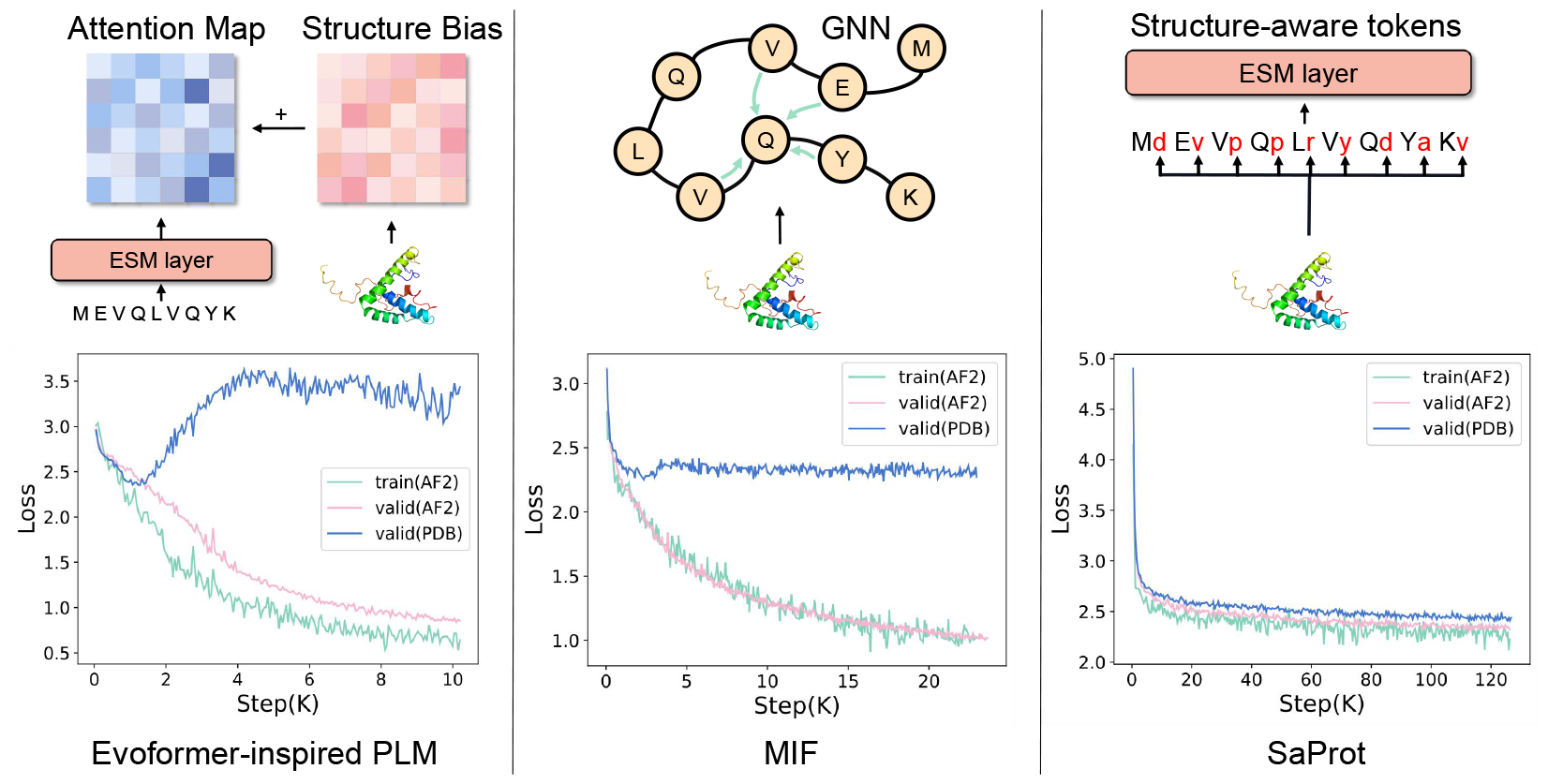
Loss trends for three protein structure models. The training set is AF2 structures while in the validation set, one is AF2 structures and the other comprises real structures from PDB.

### 3.2 Structure-aware Vocabulary

Inspired by the above discoveries, we aim to incorporate protein structures from a novel perspective. Our key idea revolves around creating a structure-aware (*SA*) vocabulary, where each *SA* token encompasses both residue and structure information, as illustrated in Figure 1.

Given a protein *P*, its primary sequence can be denoted as (*s*_1_, *s*_2_, …, *s*_*n*_), where *s*_*i*_ ∈ 𝒱 represents the residue at the *i*_*th*_ site, and 𝒱 represents residue alphabet. Building upon the concept of Foldseek, we can introduce an alternative approach for representing protein tertiary structures by using a vector quantized variational autoencoder (Van Den Oord et al., 2017). This approach enables us to develop a structure alphabet *ℱ*, wherein *P* can be represented as the (*f*_1_, *f*_2_, …, *f*_*n*_) sequence, with *f*_*j*_ ∈ *ℱ* denoting the structure token for the *j*_*th*_ residue site. To maintain simplicity, we directly adopt the default setup of Foldseek, which defines the size *m* of *ℱ* as 20.

Now, we can combine the residue and structure tokens per residue site, generating a new structure-aware sequence *P* = (*s*_1_*f*_1_, *s*_2_*f*_2_, …, *s*_*n*_*f*_*n*_), where *s*_*i*_*f*_*i*_ ∈ 𝒱×*ℱ* is the token fusing both residue and geometric conformation information. The structure-aware sequence can then be fed into a standard Transformer encoder as basic input. It’s important to note that we also introduce a mask signal “#” to both residue and structure alphabet, which results in “*s*_*i*_#” and “#*f*_*i*_” that indicate only residue or structure information is available. The size of the *SA* vocabulary is 21 × 21 = 441 (see Figure 1).

The design of this new vocabulary is simple yet innovative and fundamental, enabling the representation of any residue sequence using this “SA” sequence. As a result, protein models that utilize residue sequences as input can effortlessly integrate the new vocabulary sequence as a substitute.

### 3.3 SaProt

#### 3.3.1 Model Architecture

SaProt employs the same network architecture and parameter size as the 650M version of ESM-2. The main distinction lies in the expanded embedding layer, which encompasses 441 *SA* tokens instead of the original 20 residue tokens. This nearly identical architecture enables straightforward comparisons with the ESM model. Moreover, the model size strikes a balance between performance and feasibility for downstream task training, avoiding excessive memory or computation cost.

#### 3.3.2 Objective Function

We train SaProt using the BERT-style MLM objective, similar to ESM-1b and ESM-2, enabling the support for both protein-level and residue-level tasks. Formally, For a protein sequence *P* = (*s*_1_*f*_1_, *s*_2_*f*_2_, …, *s*_*n*_*f*_*n*_), the input and output can be represented as: *input*: (*s*_1_*f*_1_, …, #*fi*, …, *s*_*n*_*f*_*n*_) → *output*: *s*_*i*_*f*_*i*_ (see Figure 1). *fi* in #*fi* is made visible during training to reduce the model’s emphasis on predicting it. This is different from the straightforward masking strategy, i.e. randomly masking *SA* token *s*_*i*_*f*_*i*_ by “##”, and then predicting both residue and structure token directly from the *SA* vocabulary (see Appendix Figure 7). We do not adopt this strategy because the *SA* tokens may be not accurate enough^5^. Predicting the exact *SA* tokens may lead the model in the wrong optimization direction. With the proposed masking objective, although there are still inaccuracies in certain Foldseek tokens, the global structure information should remain effective, which provides valuable context for the prediction. From this perspective, it is more reasonable to predict the residue tokens rather than the Foldseek structural tokens or both of them. We perform the empirical study on the two masking strategies during pre-training in Appendix F.

To ensure a fair comparison, SaProt was pre-trained using identical training strategies with ESM-2 (refer to Appendix C). We build the pre-training dataset, which consists of approximately 40 million AF2 structures. Details are included in Appendix B, including how to proceed with the lower pLDDT region.

## 4 Experiments

We evaluate SaProt across 10 diverse downstream tasks, encompassing residue-level and protein-level tasks. Given that many proteins in the original datasets lack experimentally determined structures, we conduct all evaluations using predicted structures obtained from AlphaFoldDB without special mention. Furthermore, proteins without structures in AlphaFoldDB will not be utilized in all our experiments.

### 4.1 Zero-shot Mutational Effect Prediction

#### 4.1.1 Datasets

We adopt the ProteinGym (Notin et al., 2022) benchmark and ClinVar (Landrum et al., 2018) dataset used in Frazer et al. (2021) to evaluate the performance of SaProt on the zero-shot mutational effect prediction tasks (Meier et al., 2021). For dataset details, we refer readers to Appendix D.2.1.

#### 4.1.2 Baselines & Evaluation

We compare SaProt with two types of baselines: sequence-based models and structure-based models. For sequence-based models, we include ESM-1b (Rives et al., 2019), ESM-1v (Meier et al., 2021) (the results of 5 ESM models are averaged), ESM-2 650M (Lin et al., 2022)^6^, and Tranception L (Notin et al., 2022). For structure-based models, we consider the MIF-ST (Yang et al., 2022) and ESM-IF (Hsu et al., 2022). Additionally, we present the performance of EVE (Frazer et al., 2021), a renowed model that leverages MSA information for predicting disease variant effects, and MSA Transformer (Rao et al., 2021), a protein language model pre-trained on large scale of MSA data (we sample 384 homologous proteins for inference following Notin et al. (2022)). Here, we did not include comparisons with contrastive learning models like GearNet and 3D-PLM, as they are not directly applicable to residue-level zero-shot prediction tasks. Also note that with the exception of EVE on ProteinGym, all baseline models and their weights used in this study were obtained from the official paper. We solely employed them for prediction without any training. We trained EVE on ProteinGym ourselves using the official code as it necessitates training on each MSA.

We strictly follow the evaluation used in EVE (Frazer et al., 2021) for assessing the model’s performance on the ClinVar dataset. For the ProteinGym dataset, we employ the evaluation measures described in (Notin et al., 2022; Meier et al., 2021). Details are provided in Appendix D.2.2.

#### 4.1.3 Results

Table 1 shows the zero-shot results on ProteinGym & ClinVar, resulting in the below conclusions:

- SaProt outperforms all residue sequence-based and structure-based models on both tasks. As mentioned earlier, SaProt shares an identical network architecture, model size, and training examples with ESM-2, with the key difference lying in its structure-aware vocabulary. By comparing SaProt with ESM-2, it clear that SaProt yields consistent improvement for predicting mutational effects. Then, SaProt shows higher accuracy compared to MIF-ST, even though the latter model was trained using experimentally determined highly accurate structures.^7^ The benefit could be attributed to the large-scale structures when training SaProt. ESM-IF exhibits the poorest performance in both tasks, primarily because it was originally designed for the inverse folding task. In addition, ESM-IF model size and training data are nearly 5 times smaller than SaProt.
- MSA information enhances models’ zero-shot ability. Notin et al. (2022) introduces a technique to enhance autoregressive inference by leveraging MSA information, leading to a consistent improvement. Following it, we extend the technique to SaProt and all baselines. The results show that the integration of MSA information greatly enhances the zero-shot prediction ability of various PLMs, with SaProt still achieving the highest accuracy among them. The results also suggest that the improvement techniques used for residue sequence-based models are likely to be useful to SaProt as well.

**Table 1:** Zero-shot mutational effect prediction. ClinVar uses AUC (area under the ROC curve) and ProteinGym uses Spearman’s *ρ* as evaluation metric. They are two distinct biological tasks.

| Dataset | ESM-2 | ESM-1v | ESM-1b | Tranception L | ESM-IF | MIF-ST | EVE | MSA Transformer | SaProt |
| --- | --- | --- | --- | --- | --- | --- | --- | --- | --- |
| ClinVar | 0.862 | 0.891 | 0.900 | 0.845 | 0.748 | 0.891 | 0.878 | 0.854 | **0.909** |
| ProteinGym(w/o MSA retrieval) | 0.475 | 0.448 | 0.440 | 0.413 | 0.409 | 0.474 | - | - | **0.478** |
| ProteinGym(w/ MSA retrieval) | 0.479 | 0.472 | 0.472 | 0.465 | 0.425 | 0.480 | 0.477 | 0.464 | **0.489** |

### 4.2 Supervised Fine-tuning Tasks

#### 4.2.1 Datasets

For protein-level tasks, we evaluate SaProt on a diverse set of datasets from several benchmarks (Dallago et al., 2021; Xu et al., 2022; Rao et al., 2019), including predicting Thermostability, Metal Ion Binding, protein localization (DeepLoc), protein annotations (EC and GO) and protein-protein interaction (HumanPPI). Dataset description and splits are listed in Appendix D.3.

#### 4.2.2 Baselines

In addition to the above baselines, we compared SaProt to GearNet (Zhang et al., 2023b). Inspired by ESM-GearNet (Zhang et al., 2023a), we replaced the ESM module in ESM-GearNet with SaProt, resulting in an ensemble model called SaProt-GearNet. Training details are in Appendix D.3.3.

#### 4.2.3 Results

Experimental results are illustrated in Table 2, shedding light on the following insights:

- SaProt outperforms ESM-2 in all protein-level tasks. Specifically, SaProt shows remarkable enhancements over ESM-2 in the Thermostability, HumanPPI, Metal Ion Binding, and DeepLoc tasks. This outcome once again demonstrates that integrating structure information into PLMs leads to superior protein representation.
- SaProt outperforms the two structure models, GearNet & MIF-ST, by a substantial margin. This notable performance difference highlights the efficacy of structure modeling in SaProt.
- While SaProt outperforms the ESM models, SaProt-GearNet also outperforms ESM-GearNet, which highlights the orthogonality of SaProt with more advanced improvement techniques. However, it is interesting to note that combining two models does not always result in higher performance. For example, SaProt-GearNet and ESM-GearNet do not necessarily surpass their respective single models SaProt and ESM.

**Table 2:**
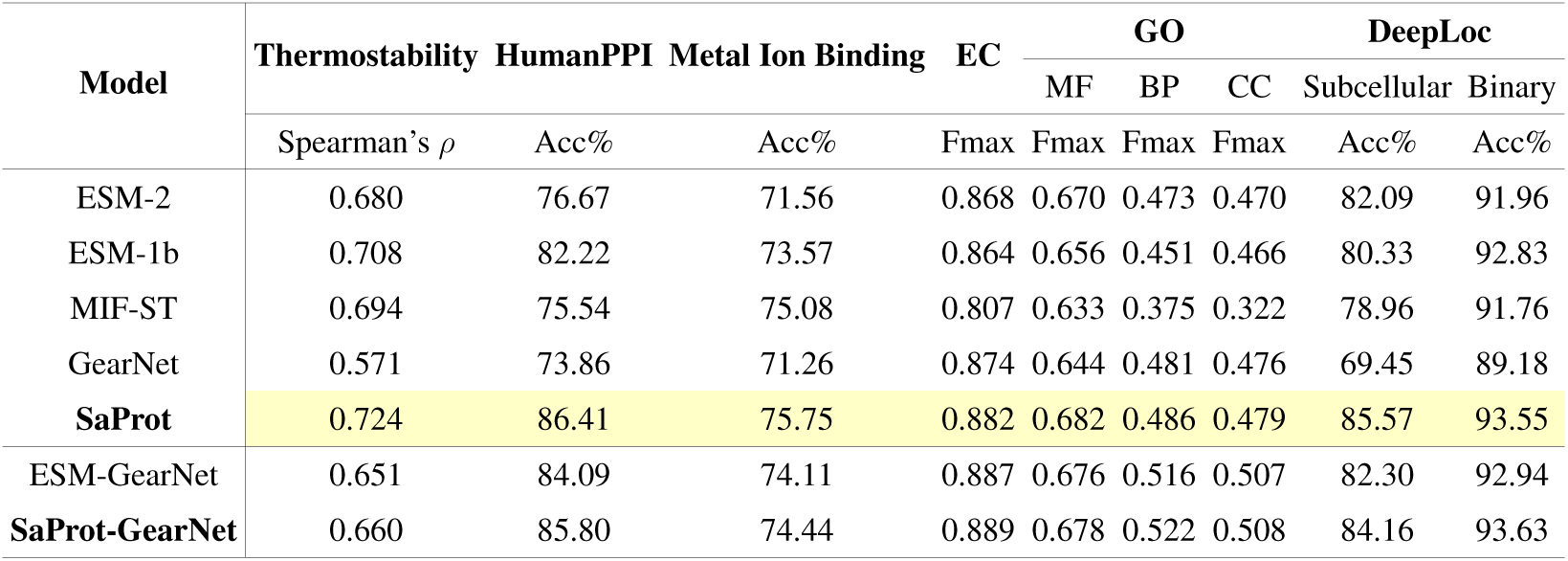
Experimental results on 8 downstream tasks.

SaProt exhibits superior performance across all tasks, when also considering the results in Section 4.1.3. Its impressive performance positions it as a compelling alternative to the ESM family.

## 5 Analysis

We conduct insightful ablation studies by dissecting SaProt. One can find more analysis in Appendix E, including comparisons of masking strategies, masking rates on structure tokens, etc.

### 5.1 Awareness Of Protein Structure

SaProt incorporates protein structure information by using structure-aware tokens rather than using explicit 3D coordinates. However, this approach relies on the accuracy of the Foldseek encoding.

Naturally, a question arises: does SaProt truly possess stronger structure information compared to ESM-2, given that residue-based PLMs also implicitly contain structure information (Rao et al., 2020)? To answer it, we conduct an additional structure prediction task, namely contact map prediction on the TAPE benchmark (Rao et al., 2019). For both SaProt & ESM-2, we freeze the backbone and solely fine-tune the contact head. The evaluation of contact map is conducted using PDB data.

As shown in Table 3, SaProt exhibits remarkable superiority over ESM-2 in the contact map prediction task, evidencing that SaProt contains more accurate structural information. From this perspective, PLM with enhanced structure feature is expected to exhibit improved accuracy in protein function prediction tasks. We additionally evaluate SaProt’s performance with all structure tokens masked as “#”, named “SaProt (Residue-only)”. SaProt (Residue-only) performs worse than ESM-2 but still exhibits a certain degree of structure prediction ability. This result demonstrates that SaProt is capable of capturing structural information even when the structure tokens are not given.

**Table 3:** Results on contact prediction. Short range, medium range and long range contacts are contacts between positions that are separated by 6 to 11, 12 to 23 and 24 or more positions, respectively.

| Model | Short Range |  |  | Medium Range |  |  | Long Range |  |  |
| --- | --- | --- | --- | --- | --- | --- | --- | --- | --- |
|  | P@L | P@L/2 | P@L/5 | P@L | P@L/2 | P@L/5 | P@L | P@L/2 | P@L/5 |
| ESM-2 | 44.87 | 44.90 | 50.18 | 45.21 | 45.80 | 53.90 | 35.33 | 41.96 | 52.11 |
| SaProt (Residue-only) | 40.29 | 40.22 | 44.10 | 36.26 | 36.69 | 42.47 | 22.15 | 27.63 | 36.68 |
| **SaProt** | **57.11** | **57.20** | **63.71** | **53.43** | **55.05** | **66.45** | **48.14** | **59.75** | **74.32** |

To further study the impact of structure and residue tokens on SaProt’s performance, we conduct an additional zero-shot prediction experiment. We randomly replace a percentage of structure tokens with random (Foldseek) tokens while keeping the residues unchanged, and then we do the opposite for residue tokens. SaProt’s performance is evaluated under this setting. As shown in Figure 3, the accuracy of SaProt decreases when either residue tokens or structure tokens are randomly substituted, which clearly emphasizes the importance of both residue and structure tokens.

**Figure 3:**
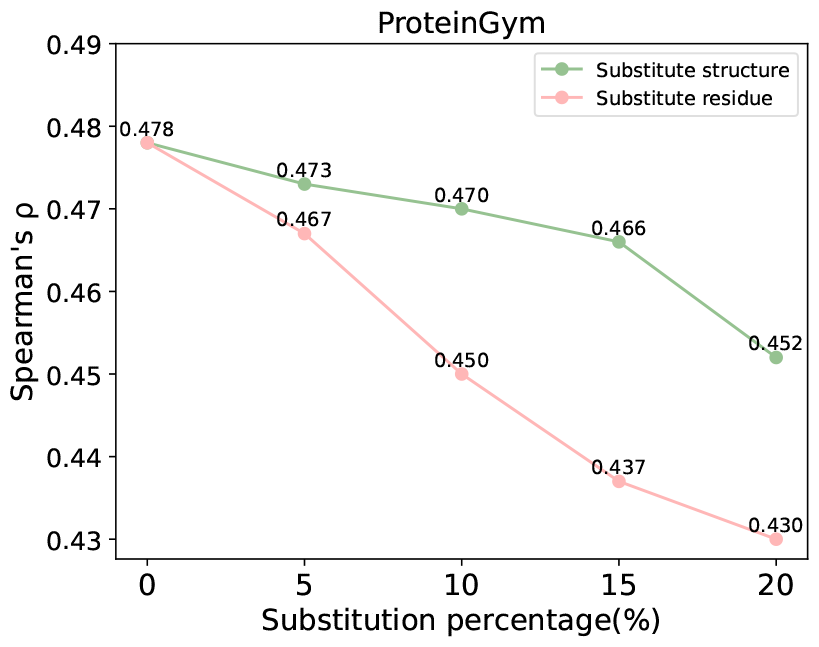
Results for different substitution percentage on (structure/residue) tokens.

### 5.2 PDB versus AlphaFoldDB

For proteins with experimentally determined structures, it is essential to investigate how SaProt performs. To do this, we continuously pre-train SaProt on 60,000 PDB structures, resulting in a variant called SaProt-PDB. We conduct evaluations by assessing SaProt and SaProt-PDB on both AF2 structures and real PDB structures. We did not evaluate all tasks due to lack of many PDB structures on some tasks.

Table 4 shows that when trained solely on AF2 structures, the overall accuracy of SaProt is not largely affected by the choice between AF2 structures or PDB structures. However, for SaProt-PDB, it is advisable to use PDB structures directly when available for downstream tasks. This may not have a substantial impact on supervised tasks such as EC and GO, as the model will be retrained on the downstream structures. However, it can have a key impact on the zero-shot task, as indicated by the comparison of 0.454 vs. 0.423 when training/testing data is highly inconsistent for SaProt-PDB.

**Table 4:** Results of SaProt and SaProt-PDB on AlphaFoldDB and PDB structures. Proteins without PDB structures on the ProteinGym dataset were removed during evaluation.

| Model | AF2 |  |  |  |  | PDB |  |  |  |  |
| --- | --- | --- | --- | --- | --- | --- | --- | --- | --- | --- |
|  | ProteinGym | EC | GO-MF | GO-BP | GO-CC | ProteinGym | EC | GO-MF | GO-BP | GO-CC |
| SaProt | **0.450** | **0.882** | **0.682** | **0.486** | **0.479** | 0.423 | 0.885 | 0.665 | 0.460 | 0.410 |
| SaProt-PDB | 0.448 | 0.880 | 0.679 | 0.482 | 0.472 | **0.454** | **0.888** | **0.669** | **0.465** | **0.415** |

In general, SaProt exhibits slightly better performance on AF2 structures, while SaProt-PDB achieves better accuracy on real structures. This outcome is expected as training and testing are consistent in the optimal performance setting. Note that some protein structures in PDB are not stored in AlphaFoldDB, so the column-level (AlphaFoldDB vs. PDB) comparison in Table 4 does not make much sense. We have released SaProt-PDB weights for utilizing PDB structures.

### 5.3 Visualization

For a more intuitive comparison, we employ t-SNE to visualize the protein representations generated by the last layer of SaProt and ESM-2. Figure 4 shows the visualization results using the non-redundant version (*PIDE <* 40%) of the SCOPe (Chandonia et al., 2018) database. For the alpha and beta proteins, the representations generated by ESM-2 are intertwined, whereas those generated by SaProt are separated based on structure type. This observation again underscores SaProt’s capability in discerning structure changes. Furthermore, we visualized the embeddings of all 400 structure-aware tokens (tokens that encompass “#” are ignored). As depicted in Figure 8 (c), we can observe a certain degree of clustering phenomenon. In the semantic space, the *SA* tokens that are in close proximity to each other often correspond to similar types of residues or Foldseek tokens.

**Figure 4:**
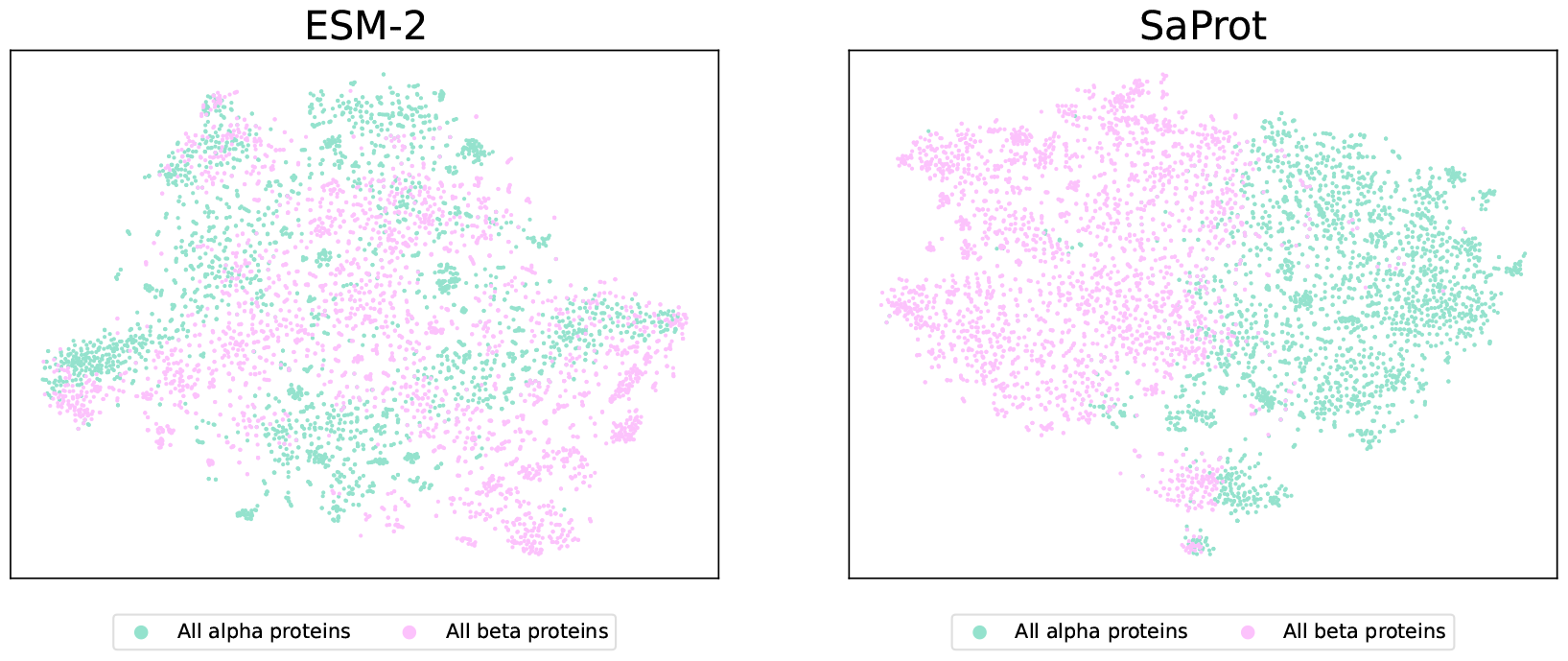
Embedding visualizations of ESM-2 and SaProt on SCOPe database.

## 6 Conclusion

In this study, we introduce a novel structure-aware (*SA*) vocabulary that integrates primary and tertiary protein structure information into the *SA*-token. This *SA*-token-based sequence has the potential to serve as a novel protein representation. Building upon this, we train a general-purpose PLM called SaProt, which achieves state-of-the-art performance on 10 protein function prediction tasks. Like the ESM models, SaProt aims to contribute to the advancement of the biological community.

This study has several limitations: (1) The performance of the proposed *SA* vocabulary heavily depends on Foldseek, which aims to balance search efficiency and encoding accuracy. Therefore, there is still room for improving the representation capability of SaProt. (2) Due to computational constraints, the model size of SaProt may not have reached its maximum capacity. (3) In addition to the mentioned tasks, there are other applications that could be explored using the SA vocabulary. For instance, predicting protein complex structures by replacing two protein sequences with SA-token-based sequences could automatically incorporate single-chain structure information. In protein generation tasks, generating SA-token sequences could potentially provide stronger structure constraints during the generation process. These avenues remain open for future research.

## Acknowledgments

We thank Sergey Ovchinnikov for many insightful suggestions for the research. Sergey advised to evaluate the performance of our model and ESM-2 on the contact map predicion task to evidence our model’s effectiveness in capuring more accurate structure information. Sergey also gave many suggestions for paper writing, figure presentation and research focus. His contributions are significant and should also be listed as an co-author. Due to abstract submission due, we cannot add his name as an author.

This work is supported by the National Key Research and Development Program of China (No. 2022ZD0115100), the National Natural Science Foundation of China (No. U21A20427), the Westlake Center of Synthetic Biology and Integrated Bioengineering (WE-SynBio), and the Research Center for lndustries of the Future (No.WU2022C030)

## A Comparison With ProstT5

A very recent preprint in Heinzinger et al. (2023) proposed ProstT5 which pre-trains the protein language model (PLM) on a mixture of data of Foldseek token sequences and residue sequences. However, they mainly focused on the bi-lingual translation between Foldseek sequence and residue sequence. ProstT5 utilizes the protein residue sequence to predict the corresponding Foldseek structure tokens, and meanwhile, it also predicts the protein residue sequence given the Foldseek structure tokens.

In contrast to ProstT5, we introduce a novel vocabulary that combines residue and Foldseek tokens, enabling the transformation of the residue sequence into an SA-token sequence. SaProt is trained on these new token sequences, effectively incorporating structure information. One acknowledged drawback of ProstT5, as stated by the original paper, is its limited ability as a general-purpose PLM, as it exhibits inferior performance in certain protein function prediction tasks.

We conducted several experiments to compare ProstT5 to SaProt, as shown in Table 5. The experimental results show that SaProt consistently outperforms ProstT5 in these protein understanding tasks, and specifically ProstT5 fails in zero-shot mutational effect prediction task.

**Table 5:** Comparison of performance between ProstT5 and SaProt.

| Model | ClinVar | ProteinGym(w/o MSA retrieval) | DeepLoc |  |
| --- | --- | --- | --- | --- |
|  |  |  | Subcellular | Binary |
| | AUC | Spearman’s $\rho$ | Acc% | Acc% |
| ProstT5 | 0.620 | 0.155 | 81.36 | 92.84 |
| **SaProt** | **0.909** | **0.478** | **85.57** | **93.55** |

## B Pre-training Data Processing

We adhere to the procedures outlined in ESM-2 Lin et al. (2022) to generate filtered sequence data, and then we retrieve all AF2 structures via the AlphaFoldDB website https://alphafold.ebi.ac.uk/ based on the UniProt ids of protein sequences, collecting approximately 40 million structures. By employing Foldseek, we encode all structures into the so-called 3Di tokens and proceed to formulate structure-aware sequences by combining residue and 3Di tokens at each position.

The AF2 structures in our study are accompanied by confidence scores, referred to as pLDDT, which provide an assessment of the precision of atom coordinates. These scores can be utilized to identify and filter out regions with low accuracy. During the MLM (masked language modeling) pretraining process, if regions with lower pLDDT scores (*<*70 as the threshold throughout this paper) are selected for the MLM prediction, we will predict the “*s*_*i*_#” token, where its input position is masked using the “##” token (note that both “##” and “*s*_*i*_#” are one token, see Figure 1). By doing so, models are forced to concentrate on predicting residue types in such regions. If regions with lower pLDDT scores are not selected for the MLM prediction, the input will be entered with the “*s*_*i*_#” token, so that models will only use the residue context in these regions to aid the prediction of other tokens. For the prediction phase of downstream tasks, we maintain complete consistency with the training data by handling the plDDT regions. Specifically, tokens within these lower pLDDT regions are represented as “*s*_*i*_#”, with only residue token visible.

## C Pre-training Details

Following ESM-2 and BERT, during training, 15% of the *SA* tokens in each batch are masked. We replace the *SA* token *s*_*i*_*f*_*i*_ with the #*f*_*i*_ token 80% of the time, while 10% of the tokens are replaced with randomly selected tokens, and the other 10% tokens remain unchanged. For the optimization of SaProt, we adopt similar hyper-parameters to those employed in the ESM-2 training phase. Specifically, we employ the AdamW optimizer (Loshchilov & Hutter, 2017), setting *β*_1_ = 0.9, *β*_2_ = 0.98 and we utilize *L*_2_ weight decay of 0.01. We gradually increase the learning rate from 0 to 4e-4 over the first 2000 steps and linearly lower it to 5e-4 from 150K steps to 1.5M steps. The overall training phase lasts approximately 3M steps. To deal with long sequences, we truncate them to a maximum of 1024 tokens, and our batch size consists of 512 sequences. Additionally, we also employ mixed precision training to train SaProt.

## D Experiments

### D.1 Data Collection

After the release of the AlphaFoldDB (Varadi et al., 2021), the majority of predicted structures are now accessible through searching UniProt IDs. We record the UniProt IDs of all proteins and query the AlphaFoldDB to retrieve all available predicted structures. On the other hand, several tasks need experimentally determined structures as training data (e.g. EC, GO and Metal Ion Binding). For these proteins, we map the PDB and chain IDs of proteins to their corresponding UniPort IDs (e.g. the chain “C” of PDB ID “6D56” will be mapped to UniPort ID “P01112”). It is worth noting that different PDB and chain IDs may correspond to the same UniProt ID (e.g. both “A” and “B” chain of PDB ID “6MAF” point to the UniProt ID “Q5D6Y5”), and they typically represent different segments within the protein. Therefore, we truncate the AF2 structures to match the corresponding PDB structures.

### D.2 Zero-shot Prediction

#### D.2.1 Datasets

##### ProteinGym

(Notin et al., 2022) is an extensive set of Deep Mutational Scanning(DMS) assays, enabling thorough comparison among zero-shot predictors. Specifically, we utilize the substitution branch, filtering out proteins with lengths exceeding 1024 or those without available structures in AlphaFoldDB. We download all AF2 structures based on UniProt ids. For evaluation, We adopt Spearman’s rank correlation as our metric.

##### ClinVar

serves as a freely accessible and publicly available repository containing information about human genetic variants and interpretations of their significance to disease (Landrum et al., 2018). In our analysis, we harness the data sourced from EVE (Frazer et al., 2021), additionally filtering out proteins with length greater than 1024 or absent from the AlphaFoldDB. To enhance the reliability of our data, we opt to consider proteins with labels 1 “Gold Stars” or higher, which indicate higher credibility. Following the methodology employed in EVE, we evaluate models’ performance using the AUC metric.

For each mutation dataset, we provide all variants with the wild-type structure, as AF2 cannot differentiate the structural changes caused by single mutations. For proteins without predicted structures in AlphaFoldDB, we simply remove them during evaluation. This also applies to all supervised task (see Table 7)

#### D.2.2 Formula

Previous residue-based PLMs like the ESM models predict mutational effects using the log odds ratio at the mutated position. The calculation can be formalized as follows:

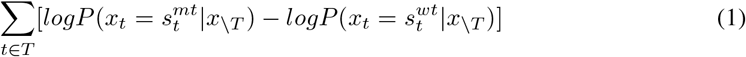

Here *T* represents all mutations and *s*_*t*_ ∈ 𝒱 is the residue type for mutant and wild-type sequence. We slightly modify the formula above to adapt to the structure-aware vocabulary, where the probability assigned to each residue corresponds to the summation of tokens encompassing that specific residue type, as shown below:

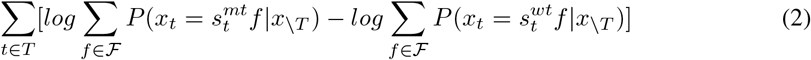

Here *f* ∈ *ℱ* is the structure token generated by Foldseek and *s*_*t*_*f* ∈ 𝒱 × *ℱ* is the structure-aware token in our new vocabulary.

#### D.2.3 Additional Comparison

We conducted additional experiments to compare SaProt to the ESM-2 15B version. Our evaluation focused exclusively on zero-shot prediction tasks, given the GPU memory constraints associated with fine-tuning the ESM-2 15B model on these supervised downstream tasks. As shown in Table 6, (1) SaProt outperformed ESM-2 15B on all zero-shot prediction tasks,(2) the larger model is not always better by comparing ESM-2 15B with ESM-2 650M. One possible reason for this could be that excessively large models may lead to overfitting issues

**Table 6:**
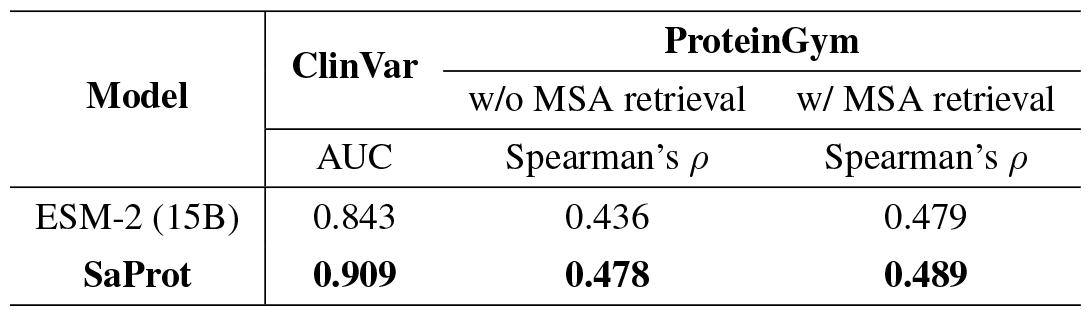
Zero-shot comparison with ESM-2 15B version. Clearly, ESM-2 15B does not improve its 650M version.

### D.3 Supervised Fine-tuning

#### D.3.1 Datasets

##### Protein Function Prediction

We compile a set of tasks that predict functions of proteins. Specifically, We employ the “Human-cell” splits of the Thermostability task from FLIP (Dallago et al., 2021), which predicts the thermostability value of proteins. Additionally, we utilize the Metal Ion Binding task (Hu et al., 2022), which is designed to predict the presence of metal ion–binding sites within a protein.

##### Protein Localization Prediction

We employ the DeepLoc (Almagro Armenteros et al., 2017) dataset to predict the subcellular locations of proteins. DeepLoc comprises two branches for sub-cellular localization prediction: one involving 10 location categories, and the other involving binary localization prediction with 2 location categories. We adhere to the original data splits.

##### Protein Annotation Prediction

We make use of two established benchmarks introduced by Deep-FRI (Gligorijević et al., 2021) to predict protein annotations encompassing multiple functional labels, i.e. Enzyme Commission(EC) number prediction and Gene Ontology(GO) term prediction. For the GO benchmark, we incorporate all three branches: Molecular Function (MF), Biological Process (BP), and Cellular Component (CC).

##### Protein-Protein Interaction Prediction

Protein-protein interaction (PPI) prediction has great potential for wide application prospects. Here we employ HumanPPI(Pan et al., 2010) from PEER(Xu et al., 2022) benchmark to predict whether two proteins interact or not.

##### Mutational effect Prediction

We employ the Fluorescence prediction and Stability prediction tasks from the TAPE (Rao et al., 2019) benchmark, the AAV dataset from the FLIP (Dallago et al., 2021) benchmark and *β*-lactamase landscape prediction from the PEER (Xu et al., 2022) benchmark. These datasets encompass mutants derived from wild-type proteins, signifying the absence of available structures.

##### Protein Structure Prediction

We adopt the contact prediction task from TAPE (Rao et al., 2019) to investigate SaProt’s awareness to protein structure.

#### D.3.2 Dataset Split

With the exception of the Metal Ion Binding and DeepLoc tasks, we utilize the official data split in the related benchmark literature (TAPE (Rao et al., 2019), PEER (Xu et al., 2022) and FLIP (Dallago et al., 2021)), which includes separate training, validation, and testing sets. Identity clustering and filtering was conducted on these benchmark datasets. For the Metal Ion Binding dataset, we perform clustering and split the data into training, validation, and testing sets based on 30% sequence identity. Note that the original dataset used in Hu et al. (2022) did not have identity clustering on the training and test sets. As the DeepLoc dataset was already clustered by 30% sequence identity, we randomly split out 20% samples from the training set as the validation set.

We summarize the dataset details in Table 7.

**Table 7:** Downstream dataset descriptions after all data pre-processing. **Category** represents a specific branch of the dataset. Note that proteins whose structures were not found in AlphaFoldDB have been removed for all baseline models during both training and testing evaluation.

| Dataset | Category | Evaluation Metric | Train | Valid | Test |
| --- | --- | --- | --- | --- | --- |
| **Protein Function Prediction** |  |  |  |  |  |
| Thermostability (Dallago et al., 2021) | Human-Cell | Spearman’s $\rho$ | 5056 | 639 | 1336 |
| Metal Ion Binding (Hu et al., 2022) | - | Acc% | 5067 | 662 | 665 |
| **Protein Localization Prediction** |  |  |  |  |  |
| DeepLoc (Almagro Armenteros et al., 2017) | Subcellular | Acc% | 8747 | 2191 | 2747 |
|  | Binary | Acc% | 5477 | 1336 | 1731 |
| **Protein Annotation Prediction** |  |  |  |  |  |
| EC (Gligorijević et al., 2021) | - | Fmax | 13089 | 1465 | 1604 |
| GO (Gligorijević et al., 2021) | BP / MF / CC | Fmax | 26224 | 2904 | 3350 |
| **Protein-Protein Interaction Prediction** |  |  |  |  |  |
| HumanPPI (Xu et al., 2022) | - | Acc% | 26319 | 234 | 180 |
| **Mutation Effect Prediction** |  |  |  |  |  |
| Fluorescence (Rao et al., 2019) | - | Spearman’s $\rho$ | 20963 | 5235 | 25517 |
| Stability (Rao et al., 2019) | - | Spearman’s $\rho$ | 53614 | 2512 | 12851 |
| AAV (Dallago et al., 2021) | 2-vs-rest | Spearman’s $\rho$ | 22246 | 2462 | 50432 |
| β-lactamase (Xu et al., 2022) | - | Spearman’s $\rho$ | 4158 | 520 | 520 |
| **Protein Structure Prediction** |  |  |  |  |  |
| Contact Prediction (Rao et al., 2019) | - | P@L | 25299 | 224 | 40 |

#### D.3.3 Training Details

In order to perform fair comparisons, we assessed our model and all baselines with the same set of hyper-parameters. we employed the AdamW optimizer, setting *β*_1_ = 0.9, *β*_2_ = 0.98 and we utilized *L*_2_ weight decay of 0.01. We consistently used a batch size of 64 and set the learning rate to 2e-5 (except 1e-3 for contact prediction). We fine-tuned all model parameters until convergence and selected the best checkpoints based on their performance on the validation set.

## E Analysis

### E.1 pre-training Comparison

#### E.1.1 Evoformer-inspired PLM

Evoformer (Jumper et al., 2021) integrates both sequence and structure information through projecting structure features as biases and incorporating them into the attention maps within the self-attention module. Nevertheless, the updates of structure features in Transformer layers are extremely time-consuming, which is infeasible to large-scale pre-training. Therefore, we simplify the interaction modules of Evoformer and employ it on standard ESM-2 model architecture. Specifically, We remove 4 triangle modules(i.e. **Triangle update using outgoing edges, Triangle update using incoming edges, Triangle self-attention around starting node** and **Triangle self-attention around ending node**) and keep the **Outer product mean** module and the **Transition** module to enable the updates of structure features. For preliminary experiments, we adopt ESM-2 35M as base model and add above modules on it to form a Evoformer-inspired PLM. We follow ProteinMPNN (Dauparas et al., 2022) to extract distance and angle features from protein structures as biases to be incorporated into the attention maps.

**Table 8:** Downstream task results for the three structure-based models. All structures used were predicted by AF2.

| Model | ClinVar | ProteinGym(w/o MSA retrieval) | DeepLoc |  |
| --- | --- | --- | --- | --- |
|  |  |  | Subcellular | Binary |
| | AUC | Spearman’s $\rho$ | Acc% | Acc% |
| Evoformer-inspired ESM(35M) | 0.589 | 0.178 | 66.11 | 89.25 |
| MIF | 0.638 | 0.256 | 61.90 | 85.90 |
| **SaProt(35M)** | **0.754** | **0.319** | **76.63** | **91.45** |

#### E.1.2 Experimental Results For Three Models

We assessed the performance of three models, i.e. Evoformer-inspired PLM ( ESM-2 35M version), MIF and SaProt 35M version, on zero-shot prediction and supervised fine-tuning tasks. The results in Table 8 are aligned with the loss change in Figure 2. For the supervised DeepLoc task, all models perform well as they are fine-tuned with new labels. Even if the pre-training is not useful, the performance after fine-tuning on new data can still be relatively good.

### E.2 Mask Rate Of Structure Tokens

PDB structures’ chains can be linked to proteins in the UniProt database, yet these structures normally constitute segments of UniProt proteins. For instance, the UniProt id “P05067” corresponds to a protein containing 770 residues, while the chain “A” within the protein of PDB id “7Y3J” has merely 110 residues from position 687 to 697 of “P05067”. Regarding AF2 structures, regions with low pLDDT values signify that certain segments of the structures lack reliability for practical use. In scenarios as described above, structures are either incomplete or not reliable, requiring models to be more robust in order to deal with such conditions.

In order to assess the robustness of SaProt, we introduced a masking procedure wherein a specific percentage of structure tokens are replaced with “#” and then we fine-tuned the model to observe resultant performance variations. Figure 5 depicts the results of fine-tuning SaProt on the DeepLoc (Almagro Armenteros et al., 2017) dataset. We employed different mask rates ranging from 0 to 1. The results show that, as the mask rate increases, there is a corresponding reduction in accuracy for SaProt. Nonetheless, the accuracy remains competitive to ESM-2. This is because even if we mask out all structural token, residual information still exists, so performance can still be recovered at least to the ESM-2 level after fine-tuning on new labels. This is different from the zero-shot prediction tasks, as discussed in Figure 3 (where performance drops significantly by substituting random structural markers). In zero-shot prediction tasks, the model does not have the opportunity to be fine-tuned, and performance is expected to drop significantly if too many structural tokens are masked, as this will lead to inconsistencies with the training data.

**Figure 5:**
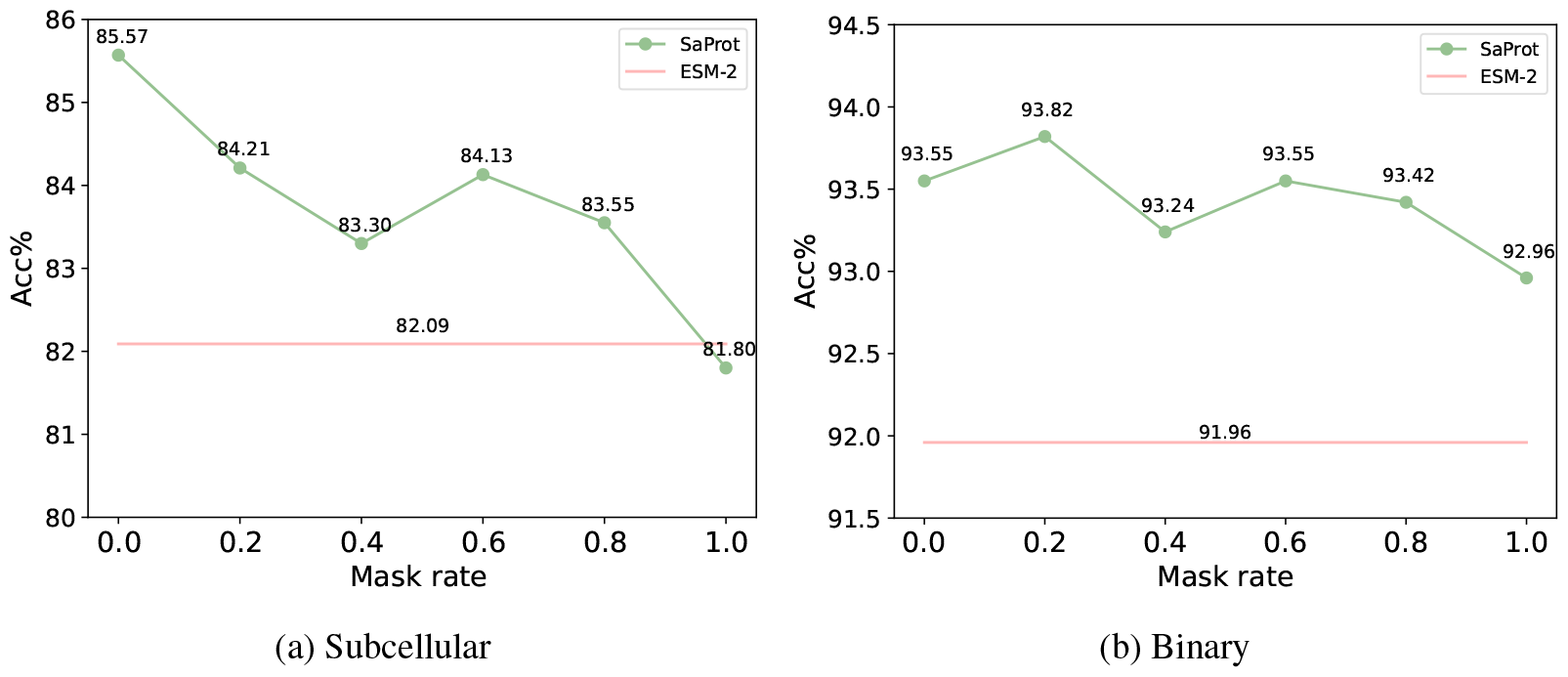
Results for various mask rates applied to structure tokens. (a) Results in the DeepLoc Subcellular branch. (b) Results in the DeepLoc Binary branch.

### E.3 Residue Sequence-only Fine-tuning

To explore the broader applications of SaProt, we evaluate its effectiveness by fine-tuning on residue sequences where all Foldseek structure tokens are substituted with “#”, resulting in “*s*_*i*_#”. This scenario often arises in various protein engineering tasks where the fitness values of protein variants can be obtained through wet experiments, but experimental structures for them are unavailable. Please note that the structures generated by AF2 may not exhibit significant distinctions for variants of the same wild-type protein.

We compare SaProt with ESM-1b and ESM-2 on four supervised mutational effect prediction datasets (Fluorescence, Stability, *β*-lactamase, and AAV). As depicted in Figure 6, SaProt performs on par with ESM-1b and ESM-2 even in the absence of structure information during the fine-tuning phase. This highlights the effectiveness of SaProt, even in situations where protein structures are not available.

**Figure 6:**
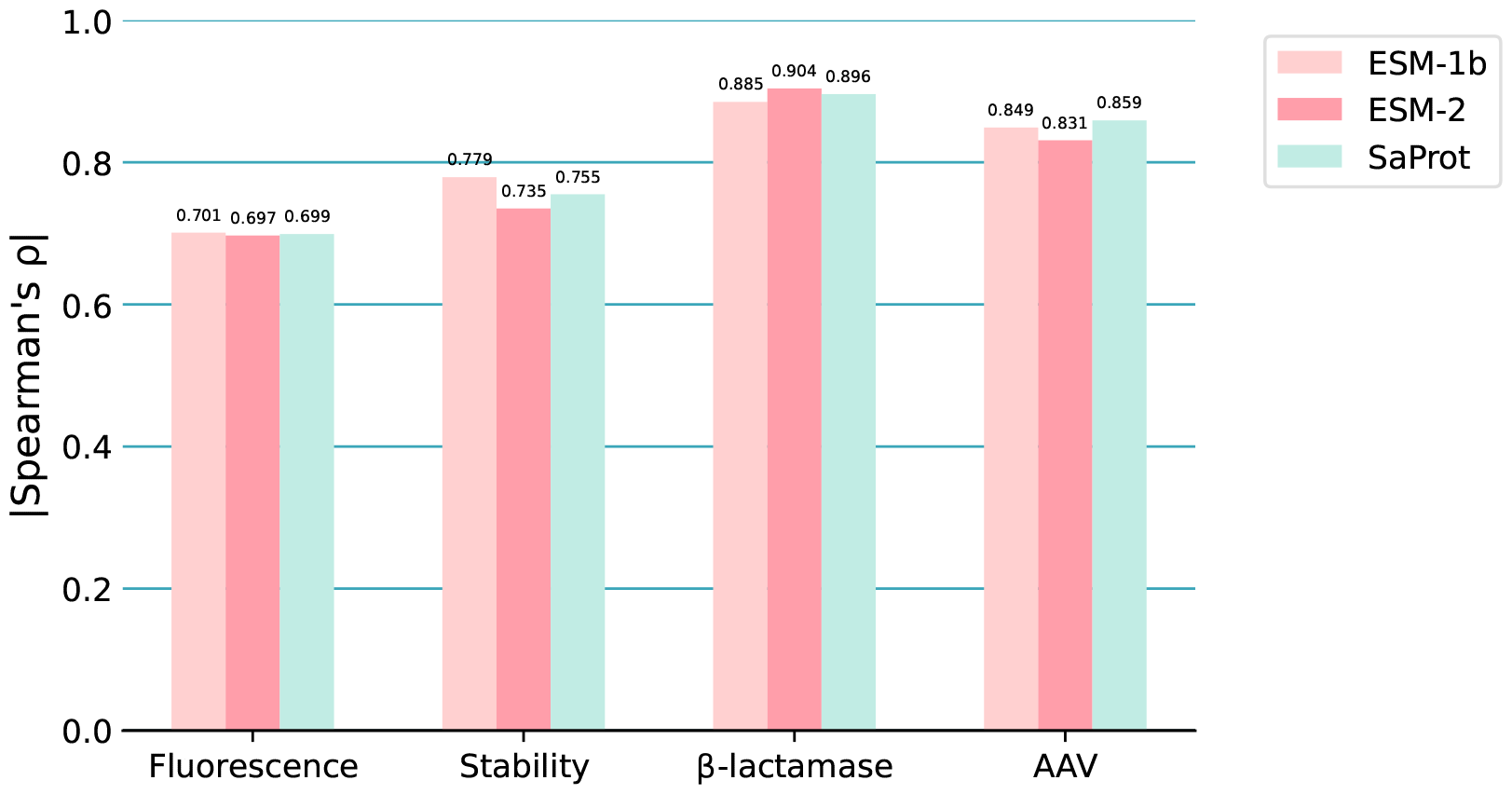
Results for residue sequence-only fine-tuning on several supervised fitness prediction datasets.

But in general, in this specific scenario, SaProt may not exhibit a clear advantage over the ESM models, but it still remains comparable in performance.

### E.4 Evaluation With ESMFold

AlphaFold2 (Jumper et al., 2021) has achieved remarkable success in predicting protein structures, with the quality of the predicted structure heavily dependent on the corresponding MSA data. However, obtaining high-quality MSA data on a large scale can be computationally intensive and time-consuming. As an alternative approach, single-sequence-based structure prediction models like ESMFold (Lin et al., 2022) can be utilized. In this study, we explore the impact of ESMFold on the performance of SaProt in both zero-shot prediction tasks and supervised fine-tuning tasks, as shown in Table 9. Our observations indicate that SaProt, utilizing structure tokens generated by ESMFold, in general achieves better or comparable accuracy to ESM-2, but falls short of its own performance with tokens generated by AF2. Hence, when feasible, it is highly recommended to employ the new vocabulary generated by AF2.

**Table 9:**
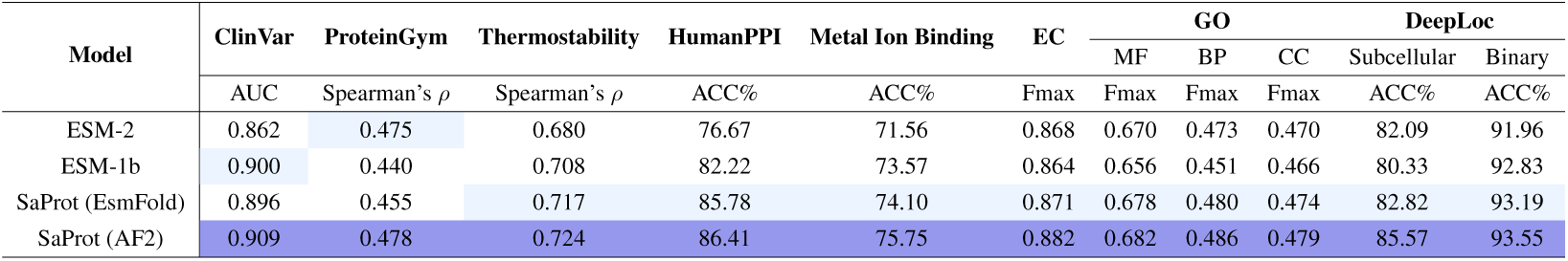
Results of SaProt using tokens generated from ESMFold and AF2.

**Figure 7:**
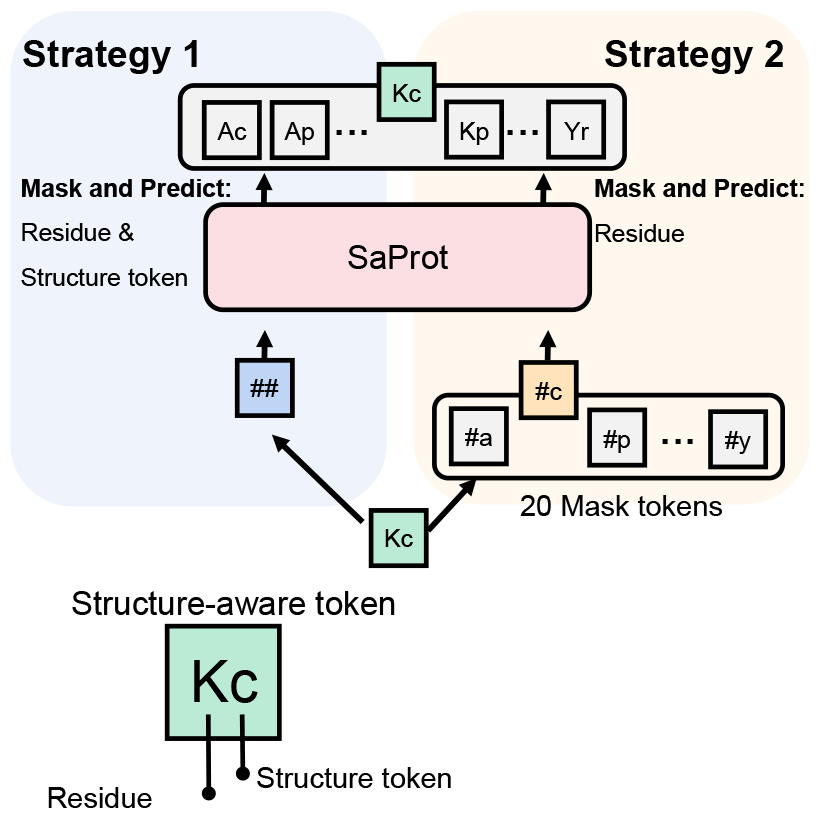
Comparison of two masking strategies.

## F Masking Strategy Comparison

For a protein structure-aware sequence *P* = (*s*_1_*f*_1_, *s*_2_*f*_2_, …, *s*_*n*_*f*_*n*_), there are two possible masking strategies that can be used, as shown in Figure 7.

### Masking Strategy 1

The most straightforward masking strategy is to randomly mask several *SA* tokens *s*_*i*_*f*_*i*_ using the symbol “##”, and subsequently predict them directly from the *SA* vocabulary. However, a potential weakness with this approach is that if the *SA* tokens are not accurate enough, predicting exact *SA* tokens may lead the model in the wrong optimization direction. This is evidenced in Table 10.

**Table 10:** Results for the two masking strategies during pre-traing phase.

| Mask&predict Strategy | ClinVar | ProteinGym |  | DeepLoc |  |
| --- | --- | --- | --- | --- | --- |
|  |  | w/o MSA retrieval | w/ MSA retrieval | Subcellular | Binary |
| | AUC | Spearman's $\rho$ | Spearman's $\rho$ | Acc% | Acc% |
| Masking Strategy 1 | 0.907 | 0.474 | 0.486 | 83.65 | 92.84 |
| Masking Strategy 2 | **0.909** | **0.478** | **0.489** | **85.57** | **93.55** |

### Masking Strategy 2

Another potential masking strategy involves either predicting the residue token *s*_*i*_ or predicting the Foldseek structure token *f*_*i*_. However, predicting *f*_*i*_ encounters the same issue mentioned above. Due to the high accuracy of residue types in protein primary sequences, predicting only the residue token seems a more effective training approach. Furthermore, predicting residue types aligns well with residue-level protein tasks, e.g., mutational effect prediction.

In Table 10, we report the results of two masking strategies on three datasets, namely ClinVar, ProteinGym, and DeepLoc. Due to highly similar results on other tasks, we omit them directly. Strategy 2 performs better as we expected, suggesting that during the training process, there might be a higher emphasis on the loss weight for predicting residues. The Foldseek structure tokens are primarily used as contextual information to aid residue prediction, rather than being utilized as labels.

## G More Visualizations

We exhibit more visualizations of learnt representations for ESM-2 and SaProt. In particular, we adopt subcellular localization and binary localization datasets, visualizing the embeddings by t-SNE (van der Maaten & Hinton, 2008). As shown in Figure 8 (a) and (b), the extracted representations from SaProt exhibit similarities or better clustering compared to those of ESM-2. Additionally, we visualized the embeddings of all 400 structure-aware tokens, as shown in Figure 8 (c). We can observe a certain degree of clustering phenomenon. In the semantic space, the *SA* tokens that are in close proximity to each other often correspond to similar types of residues or Foldseek tokens.

**Figure 8:**
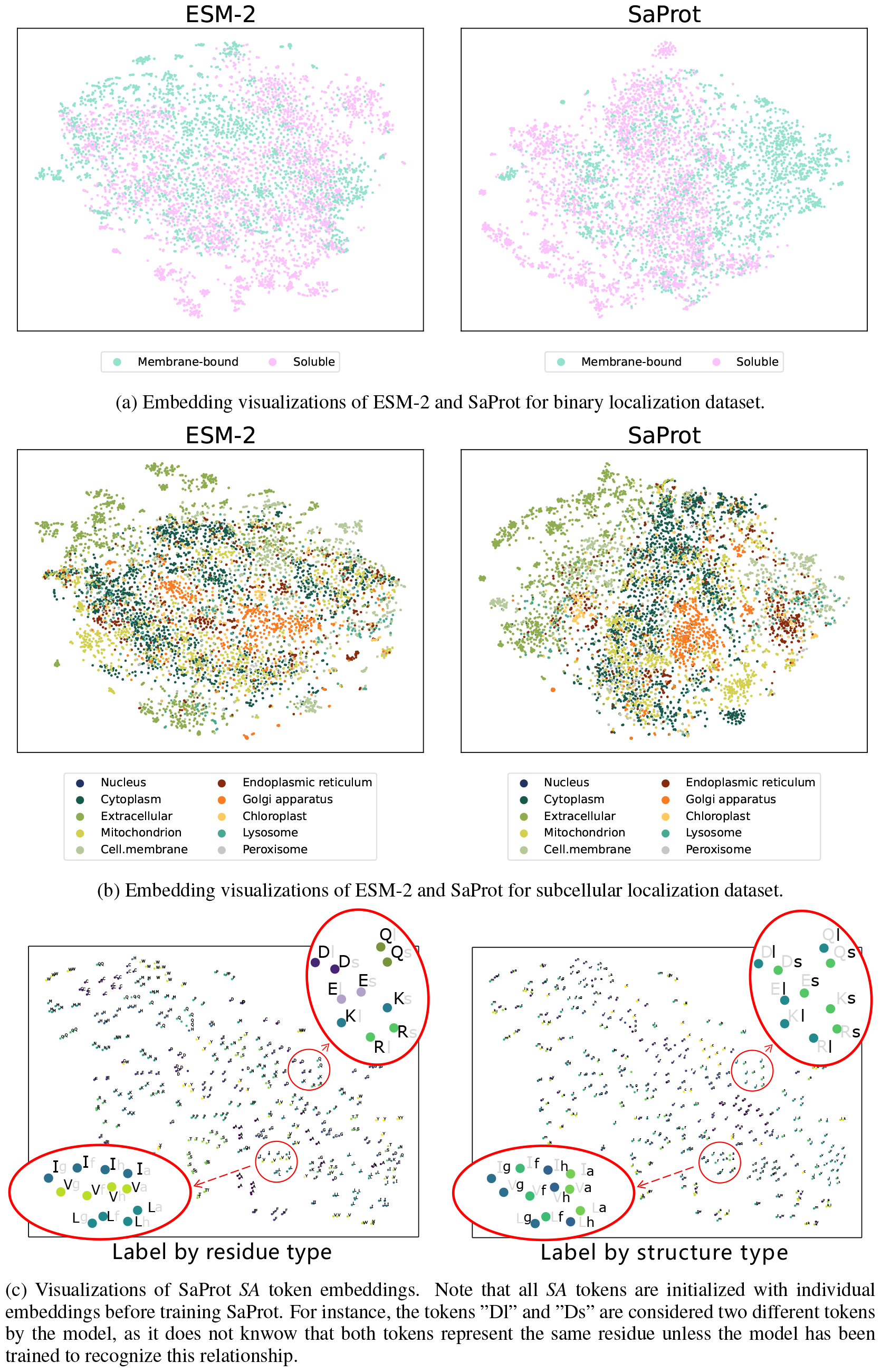
Embedding visualization

## Footnotes

1 Unlike ESM models that only offer inference code, we provide code for both training and inference.

2 Prior work employs GNNs for protein structure modeling, but GNNs suffer from the over-smoothing issue (Huang et al., 2021; Chen et al., 2020), thereby hindering large and deep protein model development.

3 Note that MIF by Yang et al. (2022) utilized only real structures for pre-training, so it is unclear whether the massive predicted structures from AF2 would be beneficial or not.

4 As a basic analysis, we utilized the 35M version of ESM-2 (see Appendix E.1.1) and SaProt. The MIF is consistent with the one described in the original paper, with a size of 3.4M.

5 The accuracy of *SA* tokens depends on the accuracy of both AF2 and Foldseek.

6 The results for 15B ESM-2 are reported in the Appendix D.2.3 which shows worse results.

7 MIF-ST exhibits poor accuracy when trained with AF2 structures, as shown in Figure 2 & Appendix E.1.2

## References

Ethan C Alley, Grigory Khimulya, Surojit Biswas, Mohammed AlQuraishi, and George M Church. Unified rational protein engineering with sequence-based deep representation learning. Nature methods, 16(12):1315–1322, 2019.

José Juan Almagro Armenteros, Casper Kaae Sønderby, Søren Kaae Sønderby, Henrik Nielsen, and Ole Winther. DeepLoc: prediction of protein subcellular localization using deep learning. Bioinformatics, 33(21):3387–3395, 07 2017. ISSN 1367-4803. doi: 10.1093/bioinformatics/btx431. URL https://doi.org/10.1093/bioinformatics/btx431.

Ivan Anishchenko, Sergey Ovchinnikov, Hetunandan Kamisetty, and David Baker. Origins of co-evolution between residues distant in protein 3d structures. Proceedings of the National Academy of Sciences, 114(34):9122–9127, 2017.

Maxwell L Bileschi, David Belanger, Drew H Bryant, Theo Sanderson, Brandon Carter, D Sculley, Alex Bateman, Mark A DePristo, and Lucy J Colwell. Using deep learning to annotate the protein universe. Nature Biotechnology, 40(6):932–937, 2022.

Tom Brown, Benjamin Mann, Nick Ryder, Melanie Subbiah, Jared D Kaplan, Prafulla Dhariwal, Arvind Neelakantan, Pranav Shyam, Girish Sastry, Amanda Askell, et al. Language models are few-shot learners. Advances in neural information processing systems, 33:1877–1901, 2020.

John-Marc Chandonia, Naomi K Fox, and Steven E Brenner. SCOPe: classification of large macro-molecular structures in the structural classification of proteins—extended database. Nucleic Acids Research, 47(D1):D475–D481, 11 2018. ISSN 0305-1048. doi: 10.1093/nar/gky1134. URL https://doi.org/10.1093/nar/gky1134.

Can Chen, Jingbo Zhou, Fan Wang, Xue Liu, and Dejing Dou. Structure-aware protein self-supervised learning, 2023.

Deli Chen, Yankai Lin, Wei Li, Peng Li, Jie Zhou, and Xu Sun. Measuring and relieving the over-smoothing problem for graph neural networks from the topological view. In Proceedings of the AAAI conference on artificial intelligence, volume 34, pp. 3438–3445, 2020.

Ratul Chowdhury, Nazim Bouatta, Surojit Biswas, Christina Floristean, Anant Kharkar, Koushik Roy, Charlotte Rochereau, Gustaf Ahdritz, Joanna Zhang, George M Church, et al. Single-sequence protein structure prediction using a language model and deep learning. Nature Biotechnology, 40(11):1617–1623, 2022.

Christian Dallago, Jody Mou, Kadina E. Johnston, Bruce J. Wittmann, Nicholas Bhattacharya, Samuel Goldman, Ali Madani, and Kevin K. Yang. Flip: Benchmark tasks in fitness landscape inference for proteins. bioRxiv, 2021. doi: 10.1101/2021.11.09.467890. URL https://www.biorxiv.org/content/early/2021/11/11/2021.11.09.467890.

Justas Dauparas, Ivan Anishchenko, Nathaniel Bennett, Hua Bai, Robert J Ragotte, Lukas F Milles, Basile IM Wicky, Alexis Courbet, Rob J de Haas, Neville Bethel, et al. Robust deep learning– based protein sequence design using proteinmpnn. Science, 378(6615):49–56, 2022.

Jacob Devlin, Ming-Wei Chang, Kenton Lee, and Kristina Toutanova. BERT: pre-training of deep bidirectional transformers for language understanding. CoRR, abs/1810.04805, 2018. URL http://arxiv.org/abs/1810.04805.

Ahmed Elnaggar, Michael Heinzinger, Christian Dallago, Ghalia Rihawi, Yu Wang, Llion Jones, Tom Gibbs, Tamas Feher, Christoph Angerer, Martin Steinegger, Debsindhu Bhowmik, and Burkhard Rost. Prottrans: Towards cracking the language of life’s code through self-supervised deep learning and high performance computing, 2021.

Jonathan Frazer, Pascal Notin, Mafalda Dias, Aidan Gomez, Joseph K. Min, Kelly Brock, Yarin Gal, and Debora S. Marks. Disease variant prediction with deep generative models of evolutionary data. Nature, 599(7883):91–95, Nov 2021. ISSN 1476-4687. doi: 10.1038/s41586-021-04043-8. URL https://doi.org/10.1038/s41586-021-04043-8.

Vladimir Gligorijević, P. Douglas Renfrew, Tomasz Kosciolek, Julia Koehler Leman, Daniel Berenberg, Tommi Vatanen, Chris Chandler, Bryn C. Taylor, Ian M. Fisk, Hera Vlamakis, Ramnik J. Xavier, Rob Knight, Kyunghyun Cho, and Richard Bonneau. Structure-based protein function prediction using graph convolutional networks. Nature Communications, 12(1):3168, May 2021. ISSN 2041-1723. doi: 10.1038/s41467-021-23303-9. URL https://10.1038/s41467-021-23303-9.

Yuzhi Guo, Jiaxiang Wu, Hehuan Ma, and Junzhou Huang. Self-supervised pre-training for protein embeddings using tertiary structures. Proceedings of the AAAI Conference on Artificial Intelligence, 36(6):6801–6809, Jun. 2022. doi: 10.1609/aaai.v36i6.20636. URL https://ojs.aaai.org/index.php/AAAI/article/view/20636.

Yan He, Xibin Zhou, Chong Chang, Ge Chen, Weikuan Liu, Geng Li, Xiaoqi Fan, Mingsun Sun, Chensi Miao, Qianyue Huang, et al. Protein language models-assisted optimization of a uracil-n-glycosylase variant enables programmable t-to-g and t-to-c base editing. Molecular Cell, 2024.

Michael Heinzinger, Ahmed Elnaggar, Yu Wang, Christian Dallago, Dmitrii Nechaev, Florian Matthes, and Burkhard Rost. Modeling aspects of the language of life through transfer-learning protein sequences. BMC bioinformatics, 20(1):1–17, 2019.

Michael Heinzinger, Konstantin Weissenow, Joaquin Gomez Sanchez, Adrian Henkel, Martin Steinegger, and Burkhard Rost. Prostt5: Bilingual language model for protein sequence and structure. bioRxiv, 2023. doi: 10.1101/2023.07.23.550085. URL https://www.biorxiv.org/content/early/2023/07/25/2023.07.23.550085.

Pedro Hermosilla and Timo Ropinski. Contrastive representation learning for 3d protein structures, 2022.

Chloe Hsu, Robert Verkuil, Jason Liu, Zeming Lin, Brian Hie, Tom Sercu, Adam Lerer, and Alexander Rives. Learning inverse folding from millions of predicted structures. ICML, 2022. doi: 10.1101/2022.04.10.487779. URL https://www.biorxiv.org/content/early/2022/04/10/2022.04.10.487779.

Mingyang Hu, Fajie Yuan, Kevin Yang, Fusong Ju, Jin Su, Hui Wang, Fei Yang, and Qiuyang Ding. Exploring evolution-aware & -free protein language models as protein function predictors. In S. Koyejo, S. Mohamed, A. Agarwal, D. Belgrave, K. Cho, and A. Oh (eds.), Advances in Neural Information Processing Systems, volume 35, pp. 38873–38884. Curran Associates, Inc., 2022. URL https://proceedings.neurips.cc/paper_files/paper/2022/file/fe066022bab2a6c6a3c57032a1623c70-Paper-Conference.pdf.

Zengfeng Huang, Shengzhong Zhang, Chong Xi, Tang Liu, and Min Zhou. Scaling up graph neural networks via graph coarsening. In Proceedings of the 27th ACM SIGKDD conference on knowledge discovery & data mining, pp. 675–684, 2021.

John Jumper, Richard Evans, Alexander Pritzel, Tim Green, Michael Figurnov, Olaf Ronneberger, Kathryn Tunyasuvunakool, Russ Bates, Augustin Žídek, Anna Potapenko, Alex Bridgland, Clemens Meyer, Simon A. A. Kohl, Andrew J. Ballard, Andrew Cowie, Bernardino Romera-Paredes, Stanislav Nikolov, Rishub Jain, Jonas Adler, Trevor Back, Stig Petersen, David Reiman, Ellen Clancy, Michal Zielinski, Martin Steinegger, Michalina Pacholska, Tamas Berghammer, Sebastian Bodenstein, David Silver, Oriol Vinyals, Andrew W. Senior, Koray Kavukcuoglu, Pushmeet Kohli, and Demis Hassabis. Highly accurate protein structure prediction with alphafold. Nature, 596(7873):583–589, Aug 2021. ISSN 1476-4687. doi: 10.1038/s41586-021-03819-2. URL https://doi.org/10.1038/s41586-021-03819-2.

Melissa J Landrum, Jennifer M Lee, Mark Benson, Garth R Brown, Chen Chao, Shanmuga Chitipiralla, Baoshan Gu, Jennifer Hart, Douglas Hoffman, Wonhee Jang, Karen Karapetyan, Kenneth Katz, Chunlei Liu, Zenith Maddipatla, Adriana Malheiro, Kurt McDaniel, Michael Ovetsky, George Riley, George Zhou, J Bradley Holmes, Brandi L Kattman, and Donna R Maglott. Clin-Var: improving access to variant interpretations and supporting evidence. Nucleic Acids Res, 46 (D1):D1062–D1067, January 2018.

Mike Lewis, Yinhan Liu, Naman Goyal, Marjan Ghazvininejad, Abdelrahman Mohamed, Omer Levy, Ves Stoyanov, and Luke Zettlemoyer. Bart: Denoising sequence-to-sequence pre-training for natural language generation, translation, and comprehension. arXiv preprint 1910.13461, 2019.

Zeming Lin, Halil Akin, Roshan Rao, Brian Hie, Zhongkai Zhu, Wenting Lu, Nikita Smetanin, Allan dos Santos Costa, Maryam Fazel-Zarandi, Tom Sercu, Sal Candido, et al. Language models of protein sequences at the scale of evolution enable accurate structure prediction. bioRxiv, 2022.

Ilya Loshchilov and Frank Hutter. Fixing weight decay regularization in adam. CoRR, abs/1711.05101, 2017. URL http://arxiv.org/abs/1711.05101.

Joshua Meier, Roshan Rao, Robert Verkuil, Jason Liu, Tom Sercu, and Alexander Rives. Language models enable zero-shot prediction of the effects of mutations on protein function. bioRxiv, 2021. doi: 10.1101/2021.07.09.450648. URL https://www.biorxiv.org/content/10.1101/2021.07.09.450648v1.

Irene MA Nooren and Janet M Thornton. Diversity of protein–protein interactions. The EMBO journal, 22(14):3486–3492, 2003.

Pascal Notin, Mafalda Dias, Jonathan Frazer, Javier Marchena-Hurtado, Aidan Gomez, Debora S. Marks, and Yarin Gal. Tranception: protein fitness prediction with autoregressive transformers and inference-time retrieval, 2022.

Xiao-Yong Pan, Ya-Nan Zhang, and Hong-Bin Shen. Large-scale prediction of human protein-protein interactions from amino acid sequence based on latent topic features. Journal of proteome research, 9(10):4992–5001, 2010.

Roshan Rao, Nicholas Bhattacharya, Neil Thomas, Yan Duan, Xi Chen, John F. Canny, Pieter Abbeel, and Yun S. Song. Evaluating protein transfer learning with TAPE. CoRR, abs/1906.08230, 2019. URL http://arxiv.org/abs/1906.08230.

Roshan Rao, Joshua Meier, Tom Sercu, Sergey Ovchinnikov, and Alexander Rives. Transformer protein language models are unsupervised structure learners. Biorxiv, pp. 2020–12, 2020.

Roshan Rao, Jason Liu, Robert Verkuil, Joshua Meier, John F. Canny, Pieter Abbeel, Tom Sercu, and Alexander Rives. Msa transformer. bioRxiv, 2021. doi: 10.1101/2021.02.12.430858. URL https://www.biorxiv.org/content/10.1101/2021.02.12.430858v1.

Alexander Rives, Joshua Meier, Tom Sercu, Siddharth Goyal, Zeming Lin, Jason Liu, Demi Guo, Myle Ott, C. Lawrence Zitnick, Jerry Ma, and Rob Fergus. Biological structure and function emerge from scaling unsupervised learning to 250 million protein sequences. PNAS, 2019. doi: 10.1101/622803. URL https://www.biorxiv.org/content/10.1101/622803v4.

Aaron Van Den Oord, Oriol Vinyals, et al. Neural discrete representation learning. Advances in neural information processing systems, 30, 2017.

Laurens van der Maaten and Geoffrey Hinton. Visualizing data using t-sne. Journal of Machine Learning Research, 9(86):2579–2605, 2008. URL http://jmlr.org/papers/v9/vandermaaten08a.html.

Michel van Kempen, Stephanie S. Kim, Charlotte Tumescheit, Milot Mirdita, Johannes Söding, and Martin Steinegger. Foldseek: fast and accurate protein structure search. bioRxiv, 2022. doi: 10.1101/2022.02.07.479398. URL https://www.biorxiv.org/content/early/2022/02/09/2022.02.07.479398.

Michel van Kempen, Stephanie S. Kim, Charlotte Tumescheit, Milot Mirdita, Jeongjae Lee, Cameron L. M. Gilchrist, Johannes Söding, and Martin Steinegger. Fast and accurate protein structure search with foldseek. Nature Biotechnology, May 2023. ISSN 1546-1696. doi: 10.1038/s41587-023-01773-0. URL https://doi.org/10.1038/s41587-023-01773-0.

Mihaly Varadi, Stephen Anyango, Mandar Deshpande, Sreenath Nair, Cindy Natassia, Galabina Yordanova, David Yuan, Oana Stroe, Gemma Wood, Agata Laydon, Augustin Žídek, Tim Green, Kathryn Tunyasuvunakool, Stig Petersen, John Jumper, Ellen Clancy, Richard Green, Ankur Vora, Mira Lutfi, Michael Figurnov, Andrew Cowie, Nicole Hobbs, Pushmeet Kohli, Gerard Kleywegt, Ewan Birney, Demis Hassabis, and Sameer Velankar. AlphaFold Protein Structure Database: massively expanding the structural coverage of protein-sequence space with high-accuracy models. Nucleic Acids Research, 50(D1):D439–D444, 11 2021. ISSN 0305-1048. doi: 10.1093/nar/gkab1061. URL https://doi.org/10.1093/nar/gkab1061.

Mihaly Varadi, Stephen Anyango, Mandar Deshpande, Sreenath Nair, Cindy Natassia, Galabina Yordanova, David Yuan, Oana Stroe, Gemma Wood, Agata Laydon, et al. Alphafold protein structure database: massively expanding the structural coverage of protein-sequence space with high-accuracy models. Nucleic acids research, 50(D1):D439–D444, 2022.

Ashish Vaswani, Noam Shazeer, Niki Parmar, Jakob Uszkoreit, Llion Jones, Aidan N. Gomez, Lukasz Kaiser, and Illia Polosukhin. Attention is all you need. CoRR, abs/1706.03762, 2017. URL http://arxiv.org/abs/1706.03762.

Jesse Vig, Ali Madani, Lav R Varshney, Caiming Xiong, Richard Socher, and Nazneen Fatema Rajani. Bertology meets biology: Interpreting attention in protein language models. arXiv preprint 2006.15222, 2020.

Fang Wu, Shuting Jin, Yinghui Jiang, Xurui Jin, Bowen Tang, Zhangming Niu, Xiangrong Liu, Qiang Zhang, Xiangxiang Zeng, and Stan Z. Li. Pre-training of equivariant graph matching networks with conformation flexibility for drug binding. Advanced Science, 9(33):2203796, oct 2022a. doi: 10.1002/advs.202203796. URL https://doi.org/10.1002%2Fadvs.202203796.

Ruidong Wu, Fan Ding, Rui Wang, Rui Shen, Xiwen Zhang, Shitong Luo, Chenpeng Su, Zuofan Wu, Qi Xie, Bonnie Berger, et al. High-resolution de novo structure prediction from primary sequence. BioRxiv, pp. 2022–07, 2022b.

Minghao Xu, Zuobai Zhang, Jiarui Lu, Zhaocheng Zhu, Yangtian Zhang, Chang Ma, Runcheng Liu, and Jian Tang. Peer: A comprehensive and multi-task benchmark for protein sequence understanding. arXiv preprint 2206.02096, 2022.

Kevin K. Yang, Niccolò Zanichelli and Hugh Yeh. Masked inverse folding with sequence transfer for protein representation learning. bioRxiv, 2022. doi: 10.1101/2022.05.25.493516. URL https://www.biorxiv.org/content/early/2022/05/28/2022.05.25.493516.

Tianhao Yu, Haiyang Cui, Jianan Canal Li, Yunan Luo, Guangde Jiang, and Huimin Zhao. Enzyme function prediction using contrastive learning. Science, 379(6639):1358–1363, 2023.

Zuobai Zhang, Minghao Xu, Vijil Chenthamarakshan, Aurelie Lozano, Payel Das, and Jian Tang. Enhancing protein language models with structure-based encoder and pre-training. In International Conference on Learning Representations Machine Learning for Drug Discovery Workshop, 2023a.

Zuobai Zhang, Minghao Xu, Arian Jamasb, Vijil Chenthamarakshan, Aurelie Lozano, Payel Das, and Jian Tang. Protein representation learning by geometric structure pretraining. In International Conference on Learning Representations, 2023b.

Gengmo Zhou, Zhifeng Gao, Qiankun Ding, Hang Zheng, Hongteng Xu, Zhewei Wei, Linfeng Zhang, and Guolin Ke. Uni-mol: A universal 3d molecular representation learning framework. In The Eleventh International Conference on Learning Representations, 2023. URL https://openreview.net/forum?id=6K2RM6wVqKu.

